# Evolutionary conservation of embryonic DNA methylome remodelling in distantly related teleost species

**DOI:** 10.1101/2023.05.24.542066

**Authors:** Samuel E. Ross, Javier Vázquez-Marín, Krista R.B. Gert, Álvaro González-Rajal, Marcel E. Dinger, Andrea Pauli, Juan Ramon Martínez-Morales, Ozren Bogdanovic

## Abstract

Methylation of cytosines in the CG context (mCG) is the most abundant DNA modification in vertebrates that plays crucial roles in cellular differentiation and identity. After fertilization, DNA methylation patterns inherited from parental gametes are remodelled into a state compatible with embryogenesis. In mammals, this is achieved through the global erasure and re-establishment of DNA methylation patterns. However, in non-mammalian vertebrates like zebrafish, no global erasure has been observed. To investigate the evolutionary conservation and divergence of DNA methylation remodelling in teleosts, we generated base resolution DNA methylome datasets of developing medaka and medaka-zebrafish hybrid embryos. In contrast to previous reports, we show that medaka display comparable DNA methylome dynamics to zebrafish with high gametic mCG levels (sperm: ∼90%; egg: ∼75%), and adoption of a paternal-like methylome during early embryogenesis, with no signs of prior DNA methylation erasure. We also demonstrate that non-canonical DNA methylation (mCH) reprogramming at TGCT tandem repeats is a conserved feature of teleost embryogenesis. Lastly, we find remarkable evolutionary conservation of DNA methylation remodelling patterns in medaka-zebrafish hybrids, indicative of compatible DNA methylation maintenance machinery in far-related teleost species. Overall, these results suggest strong evolutionary conservation of DNA methylation remodelling pathways in teleosts, which is distinct from the global DNA methylome erasure and reestablishment observed in mammals.

## BACKGROUND

During vertebrate embryonic development, the parental epigenomes are remodelled into a state that is compatible with the execution of zygotic transcriptional programs. One of the most notable examples of such epigenome reprogramming is the global erasure and re-establishment of cytosine methylation at cytosine-guanine dinucleotides (mCG) in mammalian embryos [1–6]. In mammals, the sperm genome is hypermethylated (∼80%; mCG) to a similar extent as adult somatic cells. The oocyte genome, however, contains lower levels of global DNA methylation and is characterized by large swathes of partial DNA methylation (∼40-50%; mCG) [1, 2, 4]. Upon fertilization, progressive demethylation of both parental genomes takes place until methylation is nearly completely erased by the blastocyst stage. This is followed by the re-establishment of high levels of DNA methylation coinciding with gastrulation. However, the exact molecular mechanisms underpinning early mammalian DNA methylome dynamics remain a topic of scientific debate [7–10]. Conversely, in non-mammalian vertebrates, global erasure of mCG has not been conclusively observed [11–16]. While some studies reported that DNA methylation is erased in zebrafish based on fluorescence microscopy and methyl-sensitive restriction-digestion approaches [15, 16], more robust whole-genome bisulfite sequencing (WGBS) methods have demonstrated that both the zebrafish sperm (∼90%; mCG) and oocyte (∼75%; mCG) genomes are hypermethylated.

Moreover. no global mCG erasure was observed after fertilization, with the early embryo adopting a sperm-like methylome before the onset of zygotic genome activation (ZGA) [13, 14]. Additionally, the zebrafish sperm epigenome has been shown to exist in a developmentally poised state with key developmental genes packaged into large blocks of multivalent chromatin that permit activation during ZGA [17]. Despite these differences, diverse shared epigenomic features characterize later stages of vertebrate embryonic development. These include: active DNA demethylation of enhancers during organogenesis [18-21], hypomethylation of *hox* gene clusters [13, 14, 22, 23], hypermethylation of cancer-testis antigen (CTA) gene promoters [24, 25], and accumulation of non-canonical DNA methylation (mCH; H=A,T,C) in the nervous system [26, 27].

Recently, it has been suggested that medaka (*Oryzias latipes*), a teleost (ray-finned, jaw protruding fish) like zebrafish, displays mammalian-like DNA methylome reprogrammming during early embryogenesis [28]. However, these results may have been confounded by the high content of predominantly unmethylated mitochondrial DNA present in vertebrate eggs, as this study employed an ELISA-based approach to quantify global DNA methylation levels [29]. Notably, however, medaka shares certain features with mammals that are distinct from zebrafish. For example, medaka and mammals both have distinct sex chromosomes [30], and use protamines in the packaging of their sperm genome [31,17]. It is therefore not yet clear if the global DNA methylome erasure observed during mammalian development may be evolutionarily conserved in other vertebrates, such as in medaka, or if medaka and zebrafish share teleost-specific DNA methylation remodelling features. Additionally, medaka-zebrafish hybrid embryos generated by *in vitro* fertilization using (wt) medaka sperm and zebrafish *bouncer* knockout (KO) oocytes expressing the medaka Bouncer protein, which functions as a species-specific sperm receptor, have recently been generated [32]. These embryos develop until ∼24 hpf, and transcribe RNA from both parental genomes, suggesting at least some degree of compatibility between the developmental programs in these two species [32]. These hybrids present new and exciting opportunities to study the conservation and divergence of epigenome dynamics and machinery in vertebrates.

In the current study, we have investigated DNA methylome dynamics during medaka and medaka-zebrafish hybrid embryogenesis using base-resolution whole genome bisulfite sequencing (WGBS) methodologies. Contrary to previous reports [28], we reveal that medaka embryos do not display mammalian-like remodelling of DNA methylation but instead exhibit localised DNA methylation changes comparable to those previously observed in zebrafish embryos. Medaka, like zebrafish, display hypermethylated gametic methylomes and the adoption of a paternal-like methylome state during early embryogenesis, further supporting the notion that global DNA methylome erasure is a feature, so far, only conclusively observed in mammals. Moreover, we unravel embryonic reprogramming of high levels of mCH at TGCT-containing tandem repeats in line with previous findings in zebrafish, indicative of a conserved teleost-specific phenomenon [33]. Finally, despite ∼150 million years in evolutionary divergence, we uncover remarkable conservation of both mCG and mCH DNA methylome remodelling in medaka-zebrafish hybrid embryos during the first 24 hours of embryonic development.

## RESULTS

### Comparable DNA methylome remodelling in medaka and zebrafish embryos

To quantify global 5mC levels during medaka embryogenesis, we generated base resolution DNA methylomes of developing medaka embryos at 32-cell, stage 8 early morula (st.8), stage 9 late morula (st.9), stage 10 early blastula (st.10 - ZGA), stage 11 late blastula (st.11), stage 24 - 16 somite (st.24), four days post fertilisation (4dpf), as well as of gametes and adult liver, all in biological replicates [34] (**Supplementary Table 1; Supplementary Figure S1A**). Medaka embryos exhibited DNA methylome (mCG/CG) patterning similar to what has been observed in zebrafish [13, 14] (**Supplementary Figure S1B**), with no apparent global erasure during embryogenesis (**Figure 1A**). Like zebrafish, the medaka sperm DNA methylome content (∼90%) was higher than that of the egg (∼75%), and the blastula genomes were hypermethylated like sperm (∼88%), whereas later embryonic stages (4dpf) and somatic tissues displayed lower levels of mCG (∼80%), resemblant of oocyte mCG levels (**Figure 1A**). These results are in contrast with the hypomethylated oocyte DNA methylome and the global DNA methylome erasure which has been observed in mammals [1, 2, 4], and to what has been previously reported for medaka (∼30% mCG) [28].

**Figure 1.**
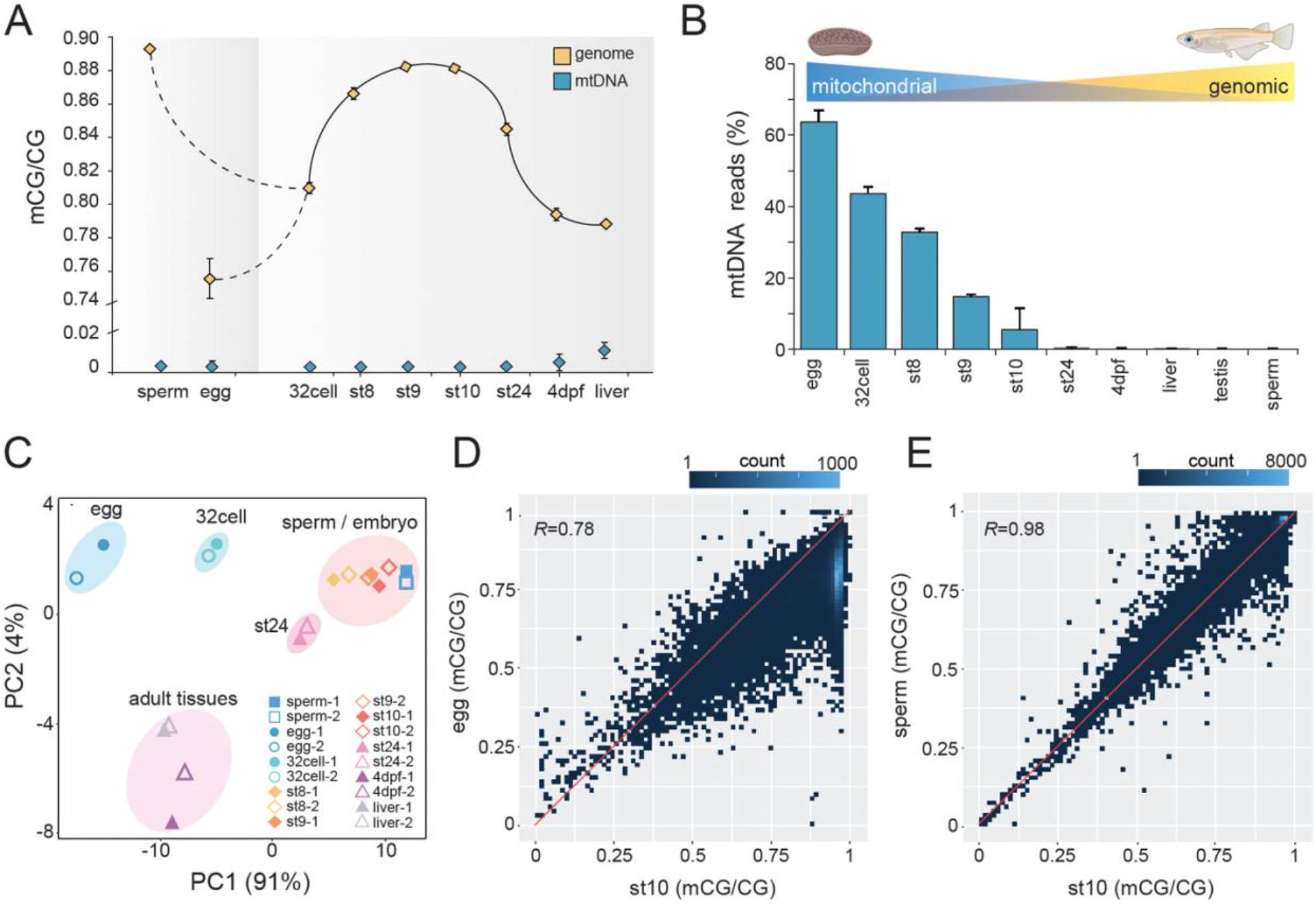
Global DNA methylation levels during medaka embryo development. **A**) Global genomic and mitochondrial DNA methylation levels (mCG/CG) in developing medaka embryos. Each point represents the mean of two WGBS replicates with error bars indicating the standard error. Stages correspond to: sperm, egg, 32 cell, stage 8 early morula (st.8), stage 9 late morula (st.9), stage 10 early blastula (st.10), stage 11 late blastula (st.11), stage 24 - 16 somite (st.24) and 4 days post-fertilisation (4dpf). **B**) Percentage of mitochondrial reads in WGBS datasets. **C)** Principal component analysis (PCA) of WGBS methylomes corresponding to developing medaka embryos (10 kb non-overlapping genomic bins). **D**) Correlation between mCG/CG (10 kb non-overlapping genomic bins) in st.10 embryos (zygotic genome activation – ZGA) and egg (right), and sperm (left). *r* indicates Pearson correlation values.

To investigate the discrepancy between our results and the previous medaka study [28], we quantified the percentage, and methylation status, of WGBS sequencing reads which mapped to the mitochondrial genome at each developmental stage (**Figure 1A, B**). As in zebrafish [13, 14], we found the mitochondrial genome to be globally hypomethylated across samples (**Figure 1A**). Furthermore, we estimate that (60 – 65%) of the DNA content in oocyte samples comes from unmethylated mitochondrial DNA (**Figure 1B**).

Such high percentage of unmethylated mitochondrial DNA in the oocyte and early embryos could explain the seemingly low levels of genomic DNA methylation reported in the ELISA-based study of medaka development [28].

To further explore the DNA methylome remodelling dynamics of medaka embryos, we performed principal component analysis (PCA) of average mCG levels in non-overlapping 10kb genomic windows across samples (**Figure 1C**). In agreement with global mCG levels (**Figure 1A**), sperm and embryonic stages clustered together, while oocytes, 32-cell stage, and somatic tissues, formed separate clusters (**Figure 1C**). Pearson correlation analysis of gametes and st.10 blastula embryos (st.10; the stage coinciding with ZGA) also revealed a high correlation between sperm and blastula methylomes (*r* = 0.98), but a lower correlation between egg and blastula methylomes (*r* = 0.78) (**Figure 1D**). Overall, our results demonstrate that medaka and zebrafish display comparable DNA methylome dynamics before ZGA and suggest that the adoption of a paternal-like methylome state before ZGA is a conserved feature of teleost development.

### Teleost-specific and pan-vertebrate epigenomic features in medaka embryos

To explore the DNA methylome of medaka development in relation to other epigenomic features, we first identified differentially methylated regions (DMRs) between adjacent developmental stages and clustered them into groups using *k*-means (*k* = 4) clustering (**Figure 2A, Supplementary Figure S2A**). We then analysed chromatin immunoprecipitation and sequencing (ChIP-seq) for key histone modifications (H3K4me3, H3K27ac, H3K27me3), and assay for transposase-accessible chromatin (ATAC-seq) datasets from medaka embryos (stage 11 and stage 24) [35–37] and compared those to the identified DMRs (**Figure 2A, Supplementary Figure S2B**).

**Figure 2.**
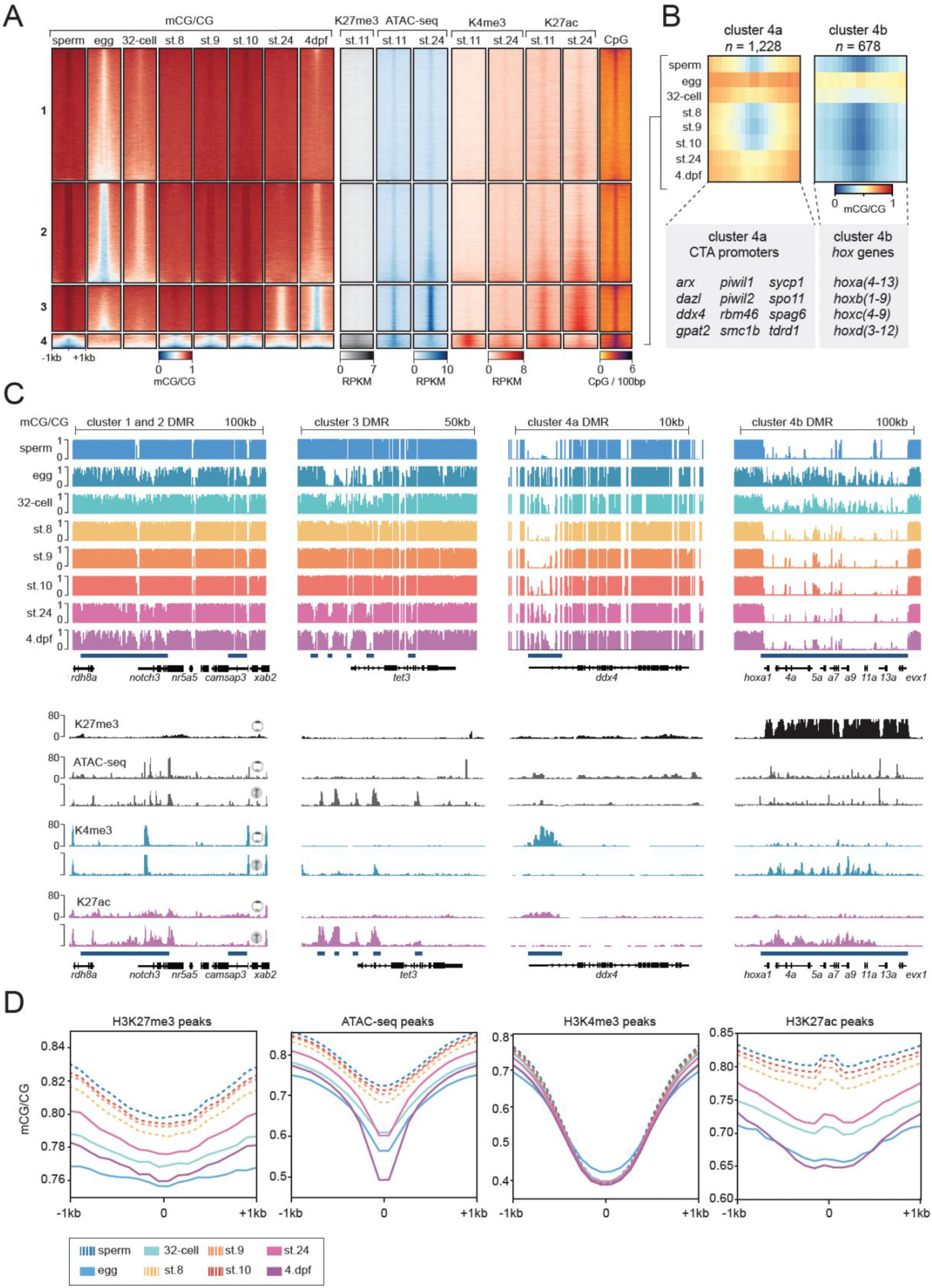
Epigenome remodelling during medaka embryo development. **A**) Heatmap of DNA methylation (mCG/CG) levels, ChIP-seq histone modification signal (H3K27me3, H3K4me3, H3K27ac) expressed as RPKM (reads per kilobase per million), ATAC-seq signal (RPKM), and CG density plotted over all developmental DMRs clustered into groups (*k*-means = 4). Embryonic stages assayed are: 32-cell, stage 8 early morula (st.8), stage 9 late morula (st.9), stage 10 early blastula (st.10), stage 11 late blastula (st.11), stage 24 - 16 somite (st.24), four days post fertilisation (4dpf), and gametes (sperm, egg). **B**) Sub-clustering of cluster-4 DMRs based on mCG/CG levels (*k*-means = 2). **C**) IGV browser snapshots of DNA methylation (mCG/CG) levels (upper panel), histone modification (H3K27me3, H3K4me3, H3K27ac), and ATAC-seq signal (RPKM) at developmental DMRs (blue boxes) during medaka development. Embryo drawings correspond to st.11 (pre-ZGA; upper drawing), and st.24 (16 somites – phylotypic period; lower drawing). Illustrations were adapted with permission from [34] **D**) Average profiles of DNA methylation (mCG/CG) during medaka embryo development plotted over H3K27me3 peaks (st.11), ATAC-seq peaks merged from st.11, st.13, st.19, st.25, st.32, H3K4me3 peaks merged from st.11, st.24) and all H3K27ac peaks identified at st.11 and st.24 that do not overlap H3K4me3 peaks. Dotted lines represent sperm and pre-ZGA/ZGA samples whereas full lines denote egg and other embryonic stages.

We found that the majority of DMRs (*n* = 54,579; clusters 1-3) are hypermethylated in the sperm and early embryo, and hypomethylated in the egg and later embryonic stages, with the three clusters mainly differing in their onset and degree of developmental demethylation (**Figure 2A, Supplementary Figure S2A**). Additionally, these three clusters mainly comprised putative enhancer regions, characterised by enrichment in ATAC-seq (open chromatin) and H3K27ac but not in H3K4me3 signal [38, 39] (**Figure 2A, Supplementary Figure S2B**). Notably, cluster 3, which displayed the earliest onset of developmental demethylation, beginning at stage 24 (phylotypic period), was also characterised by the highest enrichment in H3K27ac, and highest CpG dinucleotide density (**Figure 2A, Supplementary Figure S2A, B**). These observations are in line with previous reports that demonstrated active DNA methylation removal from CpG-rich enhancers during the organogenesis stages of vertebrate development [18, 19]. Moreover, genes associated with cluster 3 were enriched in neurodevelopmental terms, in accord with previous results in diverse vertebrate species [19] (**Supplementary Figure S2C**).

Additionally, we also identified DMRs (cluster 4; *n* =1906) with distinct dynamics to clusters 1-3, with hypermethylation in the oocyte and hypomethylation in sperm and early embryonic stages (**Figure 2A, Supplementary Figure 2A**). These regions mainly comprised promoter regions with enrichment in ATAC-seq, H3K27me3, H3K4me3 and H3K27ac, as well as high CpG density (**Figure 2A, Supplementary Figure 2B**). Further clustering of cluster 4 yielded two distinct sub-clusters (**Figure 2B**): (i) cluster 4a (*n* = 1,228), corresponding to regions hypomethylated in the sperm and early embryo but hypermethylated at later embryonic stages, such as cancer testis antigen (CTA) promoters; and (ii) cluster 4b (*n* = 678), characterized by regions which are persistently hypomethylated in the sperm and early embryo, such as the *hox* gene cluster (**Figure 2A,B Supplementary Figure S2A,B**). Representative examples of the teleost specific adoption of a paternal-like hyper-methylated genome in medaka embryos prior to ZGA (st.10), as well as examples of the conserved pan-vertebrate features of phylotypic enhancer hypomethylation, CTA promoter hypermethylation, and persistent *hox* cluster hypomethylation are visualised in **Figure 2C**, together with their corresponding changes in chromatin state.

To further assess gene-regulatory changes taking place during medaka development, we plotted mCG levels over a merged collection of enriched regions (peaks) of H3K4me3, H3K27ac and H3K27me3 data identified from st.11 and st.24 embryos as well as over ATAC-seq peaks identified from st.11, st.19, st.25, and st.32 embryos (**Figure 2D**). This analysis revealed that most promoter regions marked by H3K4me3 maintained a persistent hypomethylated state during medaka embryo development (**Figure 2D**), in line with previous reports in vertebrates [40–42]. In contrast, the majority of other regulatory regions, marked by H3K27ac, H3K27me3 and ATAC-seq signal, displayed gradual hypermethylation to match the sperm DNA methylation patterns before ZGA, followed by hypomethylation after ZGA at st.24 and 4dpf (**Figure 2D**). Overall, these results demonstrate that global hypermethylation of regulatory regions prior to ZGA, result in a sperm-like methylome in medaka, similar to what was shown previously in zebrafish [13,14,18,19,43]. Thus, these methylation dynamics are conserved in zebrafish and medaka, and potentially in all teleosts.

### Non-canonical DNA methylation dynamics in medaka embryos

Recently, we described developmental remodelling of non-canonical DNA methylation (mCH) at TGCT tetranucleotides at Mosaic Satellite Repeats (MoSat) in zebrafish and identified the teleost-specific DNA methyl-transferase - Dnmt3ba as the primary enzyme responsible for MoSat mCH deposition [33]. To interrogate whether MoSat mCH patterning, as well as global mCH dynamics, are evolutionarily conserved between zebrafish and medaka, we interrogated mCH levels in developing medaka embryos. By measuring global mCH, we observed low levels (∼0.01) of mCH at all stages of medaka embryo development (**Figure 3A**). In agreement with previous studies [33, 44], gametes (sperm, egg), displayed an increase in mCH compared to other embryonic stages (**Figure 3A**). To explore whether mCH in medaka is associated with defined sequence motifs, we identified most highly methylated mCH sites, and performed motif enrichment analysis. The topmost enriched motif displayed a similar TGCT-containing sequence to what was previously described in zebrafish (**Figure 3B**). Furthermore, the TGCT-containing motif exhibited a similar strand bias for mCH, as well as increased levels of methylation at repeats (mCH/CH > 0.1), and even more so in tandem repeats (mCH /CH > 0.25) (**Figure 3C, D**). Lastly, analysis of mCH levels across all TGCT-containing tandem repeats during medaka development revealed a reprogramming pattern nearly identical to the one previously described in zebrafish embryos (**Figure 3E, F**) (**Supplementary Figure 3**) [33]. Medaka oocytes contain relatively high levels of mCH at TGCT tandem repeats (>10%) which are then diluted during early embryo development before being re-established at comparable levels after ZGA **(Figure 3E, F)**. However, unlike in zebrafish [33], the medaka sperm did not contain significant TGCT methylation (**Supplementary Figure 3**), which could perhaps be due to incompatibility between medaka sperm protamines and this form of non-canonical mCH [31, 45] (**Figure 3E, F**). This difference notwithstanding, our findings support a conserved enrichment and remodelling of TGCT methylation in teleosts and suggest that this form of mCH could have functional roles in the regulation of ZGA.

**Figure 3.**
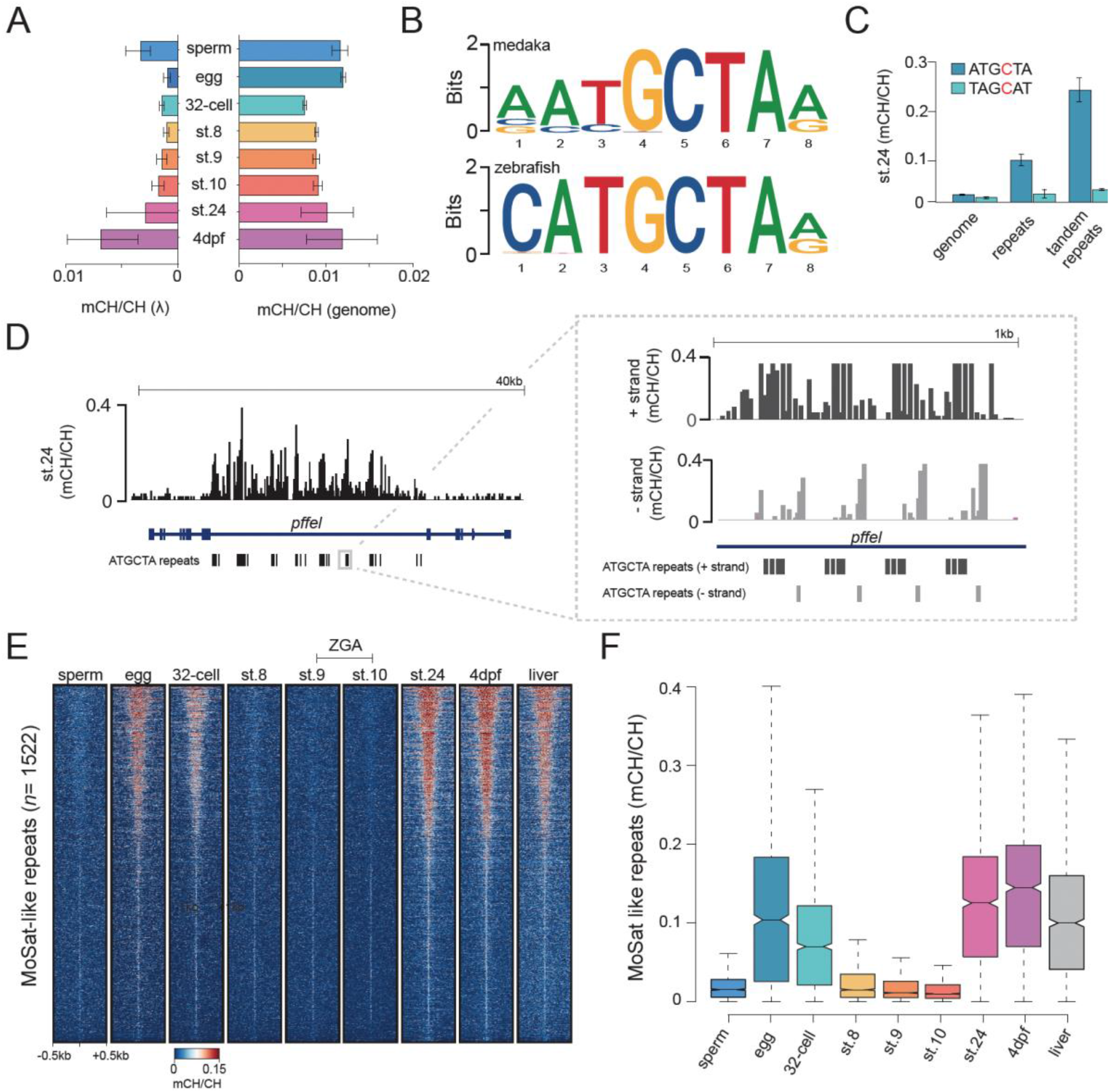
CH methylation dynamics during medaka embryo development. **A**) Global CH methylation levels (mCH/CH) in the medaka genome and in spike-in lambda (λ) controls during embryonic development. Data is represented as the mean of two WGBS biological replicates with error bars indicating the standard error. **B**) The topmost enriched motif was determined from the 10,000 most methylated CH sites in st.24 medaka embryos and 24hpf zebrafish embryos. **C**) CH methylation levels (mCH/CH) at ATGCTA and its complement TAGCAT nucleotides in the genome, repeat elements, and tandem repeat elements in st.24 medaka embryos. Data is represented as the mean of two WGBS biological replicates with error bars indicating the standard error. **D**) IGV browser snapshot of mCH/CH levels at ATGCTA repeats in st.24 medaka embryos with the right panel displaying a zoomed-in view of mCH/CH levels in a strand-specific manner. **E**) Heatmap of CH methylation levels (mCH/CH) at ATGCTA-containing tandem repeats (MoSat-like repeats), in developing medaka embryos. **F**) Distribution of CH methylation (mCH/CH) levels at ATGCTA containing tandem repeats (MoSat-like repeats) in developing medaka embryos. The box plots show the median (centre line) and the first and third quartiles (Q1 and Q3; box limits), and the whiskers extend to the last point within 1.5X the interquartile range below and above Q1 and Q3, respectively.

### Evolutionarily conserved DNA methylation patterning during medaka-zebrafish hybrid development

The generation of medaka-zebrafish (medaka ♂; zebrafish ♁) hybrids has created a powerful new model for studying parental-specific effects on epigenome regulation, as well as gene-regulatory conservation and evolution [32]. To explore the compatibility of 5mC remodelling mechanisms in these distantly related teleost species (>150 million years of divergence), we generated base resolution WGBS datasets from developing medaka-zebrafish hybrids at 3hpf, 5hpf, 8hpf and 24hpf. These time-points represent both pre- and post-ZGA periods [46, 47], for both zebrafish (ZGA at 3hpf) and medaka (ZGA between 6-8hpf) development. Additionally, 24hpf represents the start of the phylotypic period in zebrafish and the point where medaka-zebrafish hybrids become unviable [32].

Analysis of the medaka-paternal and zebrafish-maternal DNA methylome in the developing hybrids revealed persistent maintenance of the normal medaka paternal methylome, and normal adoption of the hypermethylated sperm-like methylome at the maternal zebrafish genome, up until 24 hpf (**Figure 4A**). Notably, however, these hypermethylated states persisted longer in the hybrid embryos than in normal zebrafish embryos with no loss of methylation levels by 24hpf, suggesting a potential dysregulation of DNA methylation removal from regulatory regions which normally occurs at these stages [18, 19, 21] (**Figure 4A**). Comparison of mCG levels at single CpG sites further supported strong global maintenance of the paternal medaka methylome, with high correlations (*r* = 0.98) between medaka sperm and both 3hpf and 24hpf hybrid embryos (**Supplementary Figure S4A**). Additionally, this analysis also supported the normal maternal to paternal transition of the zebrafish methylome, with high correlations (*r*= 0.98, *r*= 0.94) between *wt* and hybrid 3hpf zebrafish methylomes, and 24hpf *wt* and hybrid zebrafish methylomes, respectively (**Supplementary Figure S4A**). Furthermore, analysis of mCG/CG levels at medaka and zebrafish developmental DMRs, highlighted the establishment and maintenance of the hypermethylated sperm-like methylomes in the hybrid embryos, with an eventual lack of hypomethylation at 24hpf at key regulatory regions (**Figure 4A**). Notably, though, the paternal methylome was more affected in terms of lack of DNA methylation removal (**Figure 4B, Supplementary Figure 4B, C**). This trend could also be observed when mCG/CG levels were plotted over developmental ATAC-seq peaks (**Figure 4C)**. Accordingly, we were only able to detect a mild degree of hypermethylation at CTA gene promoters by 24hpf in both the maternal and paternal hybrid methylomes (**Figure 4D, Supplementary Figure 4B, C**). Representative examples of the normal maintenance of the global hypermethylated sperm-like methylomes, lack of hypomethylation of a subset of developmental enhancers, and mild conservation of CTA promoter hypermethylation are visualised in **Figure 4E**. Finally, plotting of mCH levels across medaka MoSat-like TGCT tandem repeats, and zebrafish MoSat tandem repeats, revealed clear increases in repeat mCH levels at both the maternal and paternal genomes in the hybrid embryos at 24 hpf, suggestive of strong conservation of the remodelling dynamics of repeat mCH [33] (**Figure 4F, Supplementary Figure 4D**).

**Figure 4.**
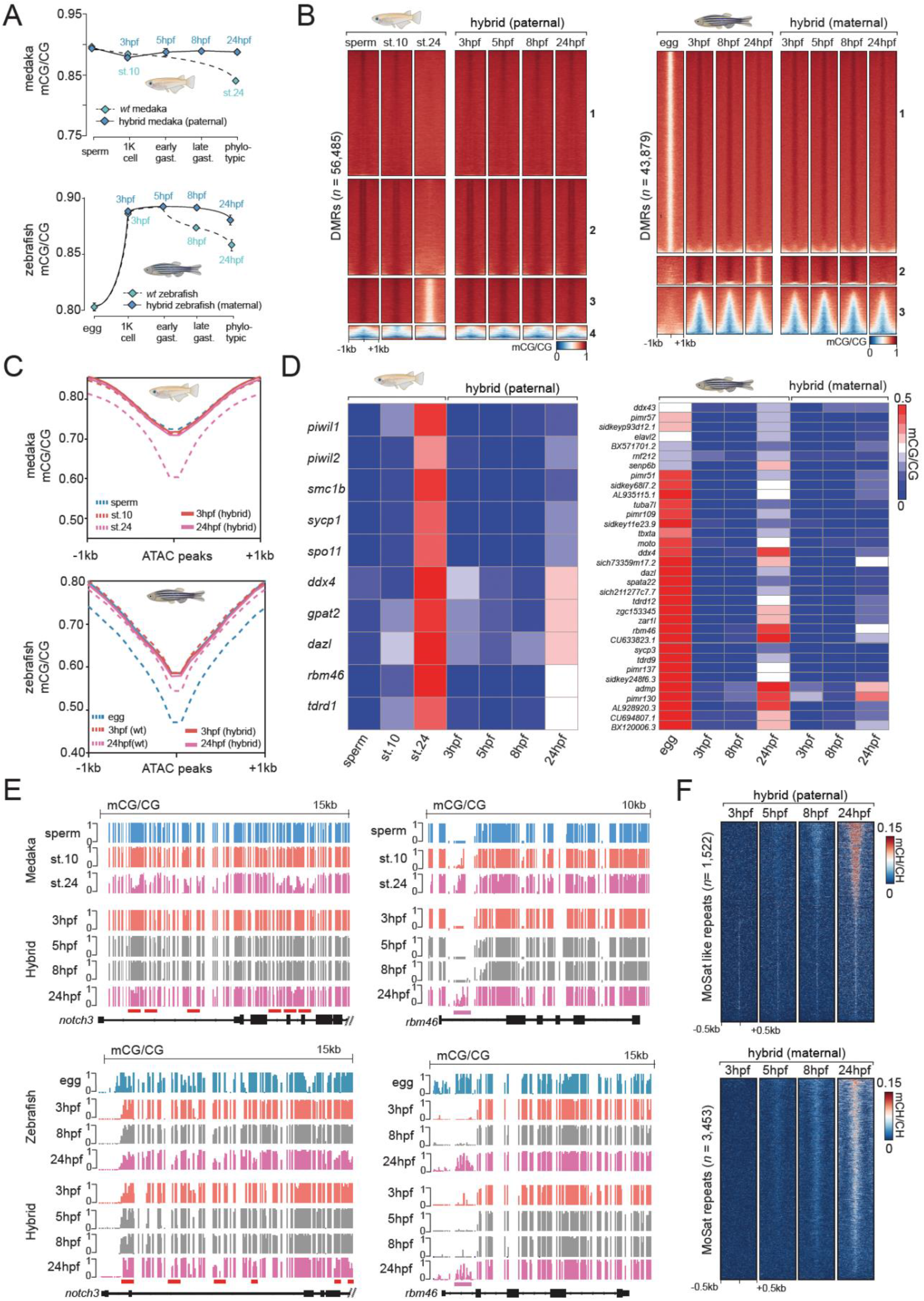
DNA methylation dynamics during medaka-zebrafish hybrid embryo development. **A**) Global mCG/CG levels of the paternal medaka methylome (top) and maternal zebrafish methylome (bottom) in medaka-zebrafish hybrids compared to the closest relative stage in wt zebrafish and medaka. Data is represented as the mean of two biological WGBS replicates with error bars indicating the standard error. **B**) Heatmap of mCG/CG levels at developmental DMRs in medaka and the paternal hybrid methylome (left), and zebrafish and the maternal hybrid methylome (right). **C**) mCG/CG plotted over developmental ATAC-seq peaks in the paternal medaka methylome (top) and maternal zebrafish methylome (bottom) from medaka-zebrafish hybrids. **D**) Heatmap of mCG/CG levels at evolutionarily conserved CTA promoters in medaka and the paternal hybrid methylome (left), and zebrafish and the maternal hybrid methylome (right). Data is represented as the mean of two WGBS biological replicates. **E**) IGV browser snapshot of mCG/CG levels during medaka, zebrafish, and medaka-zebrafish hybrid development. Red line = examples of developmental enhancers which normally undergo developmental hypomethylation. Purple line = examples of CTA promoter regions which normally undergo developmental hypermethylation. **F**) Heatmap of mCH/CH levels at MoSat-like TGCT-containing tandem repeats in the paternal medaka hybrid methylome (left) and mCH/CH levels at MoSat repeats in the maternal zebrafish hybrid methylome (right).

Overall, our results demonstrate remarkable maintenance of the medaka paternal mCG methylome and conservation of mCH dynamics in zebrafish-medaka hybrids, despite multiple cell divisions and reliance on the zebrafish cellular machinery. Additionally, we also observed uninterrupted establishment of a sperm-like hypermethylated modification state at the zebrafish maternal genome, despite the absence of the zebrafish paternal genomic contribution.

## DISCUSSION

During vertebrate embryonic development, varying degrees of epigenome remodelling take place to permit totipotency and ZGA onset. While extensive erasure and re-establishment of DNA methylation have been observed in mammals [1–6], non-mammalian vertebrates appear to require less DNA methylome remodelling during their early stages of development [13,14,48]. Additionally, no global methylome erasure has been observed in zebrafish primordial germ cells (PGCs) [24, 49], while PGCs in mammals undergo a second round of DNA methylome erasure [3, 6, 8]. In mammals, this erasure of zygotic methylomes, not only creates a state compatible with totipotency, but also removes the capacity for inheritance of DNA methylation that was parentally acquired, with notable exceptions such as the intracisternal A-particle (IAP) retrotransposons [50], and parentally imprinted regions [51]. Therefore, these findings suggest that non-mammalian vertebrates may have a higher propensity for transgenerational epigenetic inheritance when compared to mammals, as methylation acquired by the parental gametes is not erased in neither the embryo nor the developing germline of the next generation. This could mean that non-mammalian vertebrate populations could be more susceptible to factors that can influence gametic methylomes, such as environmental changes or toxins [52-54]. Additionally, this also raises the question if parentally imprinted regions, and global DNA methylome erasure are intrinsically linked, as they both occur in mammals but are absent in anamniotes.

Here we provide novel insights into DNA methylome remodelling in non-mammalian vertebrates, particularly teleosts, by generating base resolution WGBS profiles of developing medaka and medaka-zebrafish hybrid embryos. Contrary to previous reports [28], we have identified comparable DNA methylome remodelling dynamics in medaka and zebrafish. In the current study, we have found that both the medaka sperm methylome (∼90%; mCG), and oocyte methylome (∼75%; mCG), are hypermethylated and that after fertilisation the early embryo adopts a paternal-like methylome characterised by global hypermethylation of regulatory regions (**Figure 5**).

**Figure 5.**
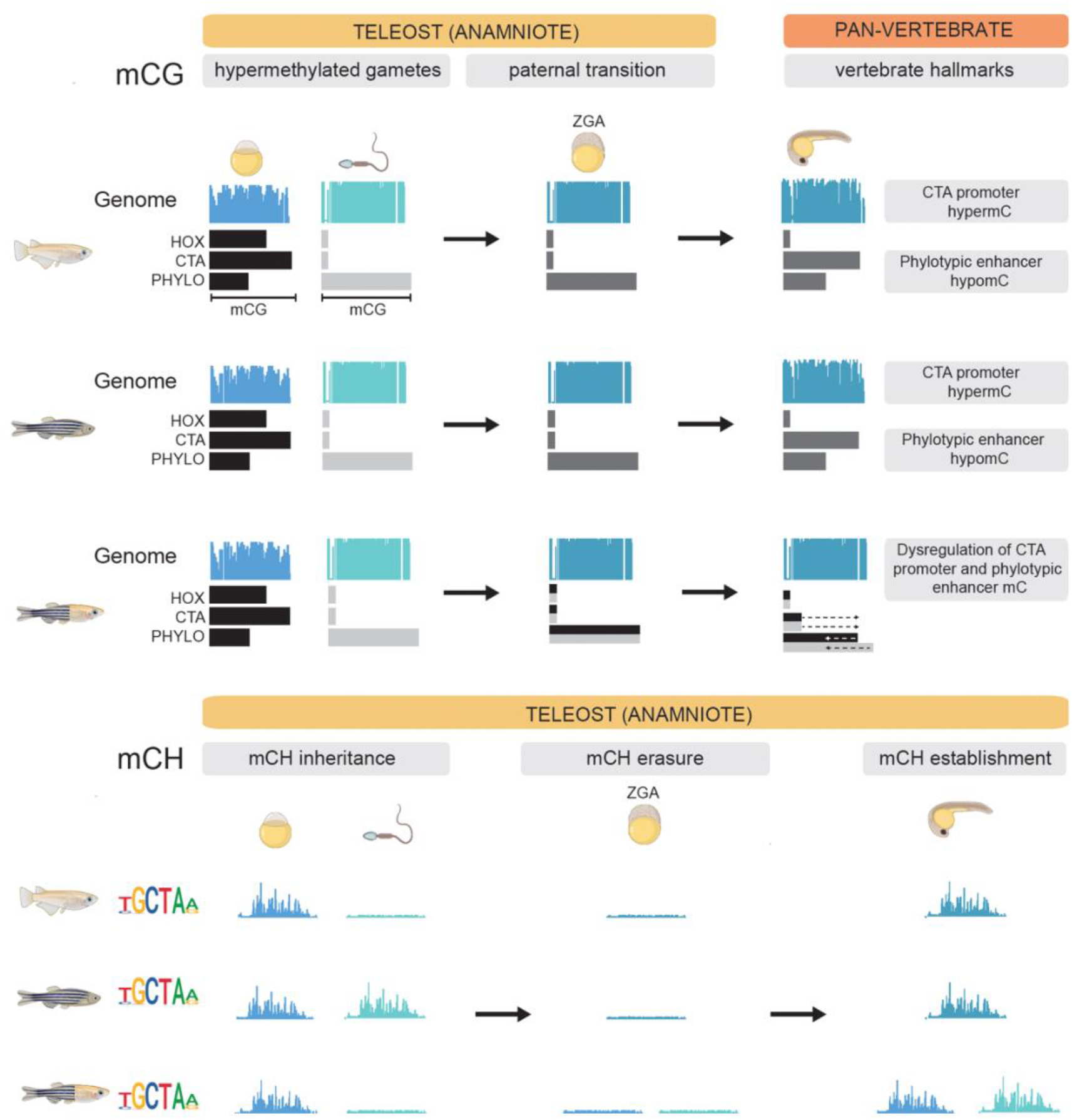
Overview of DNA methylation dynamics during teleost embryogenesis. mCG dynamics in medaka, zebrafish, and medaka-zebrafish hybrids at *hox* clusters (HOX), CTA promoters (CTA), and phylotypic enhancers (PHYLO) in gametes, ZGA blastula embryos, and during the phylotypic period (top panel). mCH dynamics at TGCT tandem repeats in medaka, zebrafish, and medaka-zebrafish hybrids in gametes, ZGA blastula embryos, and during the phylotypic period (bottom panel).

Interestingly, despite these comparable DNA methylome remodelling dynamics, medaka and zebrafish utilise different sperm packaging mechanisms; protamines and nucleosomes respectively [31, 36, 45]. It has previously been demonstrated that zebrafish sperm, and early zebrafish embryos, maintain hypomethylated regions via so called “placeholder” nucleosomes composed of H3K4me1 and the histone variant H2A.Z(FV) [43]. Whether a similar regulatory mechanism could be compatible with the protamine-rich medaka sperm, remains to be determined.

Apart from the evolutionary conservation of mCG remodelling, our analyses also reveal conservation of developmental reprogramming of high mCH levels at TGCT-containing repeats during embryogenesis (**Figure 5**). This form of mCH is distinct from mammalian CAG methylation observed in oocytes and embryonic stem cells and from the vertebrate-specific CAC methylation in neurons [26, 27, 55–57]. However, the developmental reprogramming of TGCT methylation is reminiscent of global mCH dynamics during the mammalian life cycle, as it is characterised by mCH inheritance from gametes, which is followed by its erasure coinciding with ZGA and re-establishment at later embryonic stages [8, 44]. Moreover, it was previously shown that zebrafish TGCT methylation is driven by the teleost-specific enzyme Dnmt3ba [33] and thus our findings suggest that its role in mCH deposition may be conserved across fishes. However, unlike in zebrafish, medaka sperm was depleted of mCH which could indicate a dependency of TGCT methylation on the nucleosomal genome structure (**Figure 5**). Overall, our observations further highlight an unexplored diversity of mCH systems and DNMT targets in vertebrates and suggest a possible function for mCH in ZGA regulation.

In the current study, we took advantage of the previously described medaka-zebrafish-hybrid system to study the interspecies compatibility of early DNA methylome remodelling [32]. Hybrid organisms are valuable assets in exploring the relationship between genome evolution, epigenetic control, and cytoplasmic components. For example, hybrids were used to elucidate factors controlling fertilisation and ZGA timing [32,58,59] and to explore genome evolution as exemplified by studies in *Xenopus* [60-62]. Here we have generated first base-resolution DNA methylome profiles of an inter-order hybrid organism [32] to explore the evolution and conservation of DNA remodelling pathways and machinery over large evolutionary distances. Despite >150 million years in evolutionary distance, we observe remarkable conservation of both medaka-paternal, and zebrafish-maternal DNA methylome dynamics, in both the mCG and mCH context during the first 24 hours of embryonic development. The paternal medaka DNA methylome was fully maintained in the presence of zebrafish cytoplasmic components during early development, which involves periods of rapid cell division and reliance on maternal zebrafish oocyte proteins. Moreover, the maternal zebrafish methylome was remodelled normally to match the paternal-like methylome state prior to ZGA (**Figure 5**). Importantly, these results are also in agreement with earlier UV irradiation studies which demonstrated that the zebrafish maternal-to-paternal DNA methylome transition was not dependent on the paternal methylome being present as a remodelling template [13].

This work thus supports the notion that the maternal-to-paternal methylome transition is entirely maternally driven. However, at 24hpf, when the hybrid embryos become inviable, defects in key developmental enhancer demethylation, and CTA promoter hyper-methylation were detected, particularly on the paternal medaka methylome. Normally, during the phylotypic period – the most conserved phase of vertebrate development - medaka and zebrafish display comparable epigenomic and transcriptomic profiles [37]. It is currently unclear what could be driving these DNA methylome remodelling defects, and if they are a cause or the consequence of arrested development, which characterises medaka-zebrafish hybrid embryos [32]. However, some of these defects could be explained by the insufficient production of appropriate zebrafish or medaka proteins implicated in DNA methylome remodelling events as well as the exhaustion of maternal protein pools. This could be of particular importance for cases where incompatibility exists between the target sequence of one organism and the protein specificity of the other one. Such a scenario is likely more important for the dysregulation of CTA promoters, as CTA genes, and their degree of hypermethylation, differ between species. On the other hand, developmental enhancer demethylation occurs at tens of thousands of regions, has roots in invertebrate systems [63], and is likely less sequence dependent. Furthermore, in medaka-zebrafish hybrids, ZGA onset is determined by the maternal zebrafish ZGA timing [58], which results in paternal medaka proteins being expressed earlier than they normally would. This again, could cause an imbalance in the cellular factors required for proper DNA methylome remodelling. Contrarily, we did not observe any major defects in mCH remodelling (**Figure 5**), which indicates either a compatibility between the Dnmt3ba enzymes and their targets for both species or a higher resilience to alterations in Dnmt3ba levels. Overall, this work sheds new light on the evolutionary conservation, diversity, and compatibility of DNA methylome remodelling in vertebrates, and finds no evidence for global DNA methylome erasure in teleosts, which has thus far only been conclusively described in mammalian embryos. Finally, this raises the possibility that all teleosts, and potentially all non-mammalian vertebrates, may display a higher propensity for DNA methylation-based epigenetic inheritance - which could have important ecological ramifications.

## METHODS

### Zebrafish husbandry

Zebrafish experiments were performed at the Garvan Institute of Medical Research in accordance with the Animal Ethics Committee AEC approval and with the Australian Code of Practice for Care and Use of Animals for Scientific Purposes.

### Medaka husbandry

The medaka (*Oryzias latipes*) iCab wild-type strain was maintained and embryos staged as previously described [34]. All experimental protocols have been approved by the Animal Experimentation Ethics Committees at the Pablo de Olavide University and CSIC (license number 02/04/2018/041).

### Generation of medaka-zebrafish hybrids

*In vitro* fertilization experiments for generating hybrids were conducted according to Austrian and European guidelines for animal research and approved by the Amt der Wiener Landesregierung, Magistratsabteilung 58–Wasserrecht (animal protocols GZ 342445/2016/12 and MA 58-221180-2021-16 for work with zebrafish; animal protocol GZ: 198603/2018/14 for work with medaka). Medaka-zebrafish hybrids were generated as previously described [32].

### Genomic DNA extraction

Medaka, zebrafish, and medaka-zebrafish hybrid samples were dissolved in homogenization buffer (20 mM Tris pH 8.0, 100 mM NaCl, 15 mM EDTA, 1% SDS, 5 mg/mL proteinase K) for three hours at 55°C followed by two Phenol/Chloroform/Isoamyl Alcohol (25:24:1,PCI) extractions. PCI extraction was performed using an equal volume of PCI to sample. The mixture was centrifuged for five minutes at 13,000 rpm to separate the phases. DNA was then precipitated from the aqueous phase by the addition of 1/10 volume of 3M NaOAc, 20 μg/mL linear acrylamide and three volumes of ice-cold ethanol and incubated for two hours at -20°C. DNA was pelleted by centrifugation for 30 minutes, washed with 75% ethanol, and resuspended in nuclease-free water.

### WGBS library preparation

WGBS libraries were prepared from each developmental stage in biological replicates. Unmethylated lambda phage DNA (0,5%; Promega, Madison, WI, USA) was spiked into each DNA sample before the DNA was sonicated to an average of 300bp. DNA was then purified using AMPure XP beads (Beckman Coulter, Lane Cove, NSW, Australia) and bisulfite converted using the EZ DNA Methylation-Gold Kit (Zymo Research, CA, USA), following manufacturer’s instructions. Bisulfite-converted DNA was then processed to generate WGBS libraries using the Accel-NGS™ Methyl-Seq DNA Library Kit (Swift Biosciences, Ann Arbor, Mi, USA), following manufacturer’s instructions. WGBS libraries with 15% PhiX spike-in were sequenced on the Illumina HiSeq X platform (150bp PE sequencing, high output mode).

### WGBS data processing

WGBS sequencing reads were hard-trimmed using Trimmomatic to remove the adapter sequences introduced during library preparation (ILLUMINACLIP:adapters.fa:2:30:10 SLIDING WINDOW:5:20 LEADING:3 TRAILING:3 CROP:130 HEADCROP:20) [64].

Trimmed reads were then mapped using WALT [65] onto either the bisulfite-converted *Oryzias latipes* ASM223467 or zebrafish danRer10 reference containing the λ genome as a separate chromosome. Hybrid samples were mapped onto a custom genome containing both the *Oryzias latipes* ASM223467 and zebrafish danRer10 genomes. The resulting SAM files were converted to BAM format and the percentage of reads mapping to the mitochondrial genome was calculated. The BAM files were deduplicated using sambamba markdup [66] before CG (--mergeContext) and CH methylation levels (--CHH, --CHG) were called using MethylDackel extract (https://github.com/dpryan79/MethylDackel). Genomic data was visualized in the IGV browser [67].

### DNA methylation analysis

Global methylation levels were calculated from methylDackel-generated bedGraphs by dividing the sum of all methylated cytosine calls (column 5) by the number of total cytosine calls (sum of columns 5 and 6). Principal component analysis was performed on 10kb genomic bins using the prcomp function in R and plotted with ggplot2. Scatterplots were generated with the geom_bin2d function in ggplot2 ((bins=75) + geom_smooth(method=lm). Pearson correlations were calculated with the Hmisc rcorr function in R. Differentially methylated regions were calculated using DSS (delta=0.2, p.threshold=0.05, minlen=50, minCG=5, dis.merge=100, pct.sig=0.5) [68] and methylation averages of these regions calculated with bedtools map [69]. Heatmaps were generated using deepTools computeMatrix (computeMatrix reference-point --referencePoint centre -b 1000 -a 1000 -bs 25) [70] and the NAN values in matrices were replaced with average DNA methylation levels before plotting with the plotHeatmap function and plotProfile functions (--plotType heatmap --yMin 0 --yMax 0.15 -- perGroup). mCH motif analysis was performed on top 10,000 most highly methylated mCH sites with minimal 10X coverage identified in st.24 medaka embryos. MEME software with the target sequence being the methylated site +/- 4bp, was used for motif identification [71].

### ChIP-seq and ATAC-seq analysis

ChIP-seq and ATAC-seq sequence reads were trimmed with Trimmomatic with the following settings respectively: ILLUMINACLIP:TruSeq3.fa:2:30:10 SLIDINGWINDOW:5:20 LEADING:3 TRAILING:3 MINLEN:20 and ILLUMINACLIP:Nextera.fa:2:30:10 SLIDINGWINDOW:5:20 LEADING:3 TRAILING:3 MINLEN:20 CROP:130 HEADCROP:150). Trimmed reads were then mapped with bowtie2 using default settings [72]. The resulting alignments in BAM format were deduplicated using sambamba markdup [66]. ATAC-seq BAM files were filtered to remove reads with fragment sizes greater than 100 bp and peaks from ATAC-seq data were called using MACS2 [73]. RPKM bigWigs were generated using the deepTools bamCoverage function (--normalizeUsing RPKM --centerReads) [70].

## Supporting information

Ross_et_al_2023_TableS1

## AVAILABILITY OF DATA AND

### MATERIALS

Data generated for this submission have been uploaded to ArrayExpress https://www.ebi.ac.uk/arrayexpress/ under the accession number E-MTAB-12535

## ACKNOWLEDGEMENTS

We acknowledge the Kinghorn Centre for Clinical Genomics Sequencing Laboratory for the generation of whole genome bisulfite sequencing libraries. Drawings were created with the help of BioRender.com software. The authors thank Alex de Mendoza for critical reading of the manuscript.

## FUNDING

The Australian Research Council (ARC) Discovery Project (DP190103852); Ramón y Cajal fellowship (RYC2020-028685-I) and the “Proyecto de Generación de Conocimiento 2021” project (PID2021-128358NA-I00) from the Spanish Ministry of Science and Innovation, as well as funding from CEX2020-00108-M Unidad de Excelencia María de Maeztu to O.B. supported this work. KRBG was supported by a DOC Fellowship from the Austrian Academy of Sciences. Work in the Pauli lab was supported by the FWF START program (Y 1031-B28 to AP), the ERC CoG 101044495/GaMe, the HFSP Career Development Award (CDA00066/2015 to AP), a HFSP Young Investigator Award (RGY0079/2020 to AP) and the FWF SFB RNA-Deco (project number F80). The IMP receives institutional funding from Boehringer Ingelheim and the Austrian Research Promotion Agency (Headquarter grant FFG-852936). For the purpose of Open Access, the author has applied a CC BY public copyright license to any Author Accepted Manuscript (AAM) version arising from this submission.

## AUTHOR INFORMATION

OB conceived the study. SR performed bioinformatics analyses. JVM and JRMM generated and collected the medaka samples. Medaka-zebrafish hybrid samples originated from AP’s lab and were generated and collected by KRBG. AGR, MD and OB participated in data analysis. SR and OB wrote the manuscript. All authors contributed to, read, and approved the final manuscript.

## ETHICS DECLARATION

The animal study was reviewed and approved by the Garvan Institute of Medical Research Animal Ethics Committee (AEC approval 20/09), Animal Experimentation Ethics Committees at the Pablo de Olavide University and CSIC (license number 02/04/2018/041), and by the Amt der Wiener Landesregierung, Magistratsabteilung 58-Wasserrecht (animal protocols GZ 342445/2016/12 and MA 58-221180-2021-16 for work with zebrafish; animal protocol GZ: 198603/2018/14 for work with medaka).

## COMPETING INTERESTS

The authors declare that they have no competing interests.

## Notes

https://www.ebi.ac.uk/arrayexpress/

### Competing Interest Statement

The authors have declared no competing interest.

