## Supplementary_Figures_S1-S4 for "Evolutionary conservation of embryonic DNA methylome remodelling in distantly related teleost species"

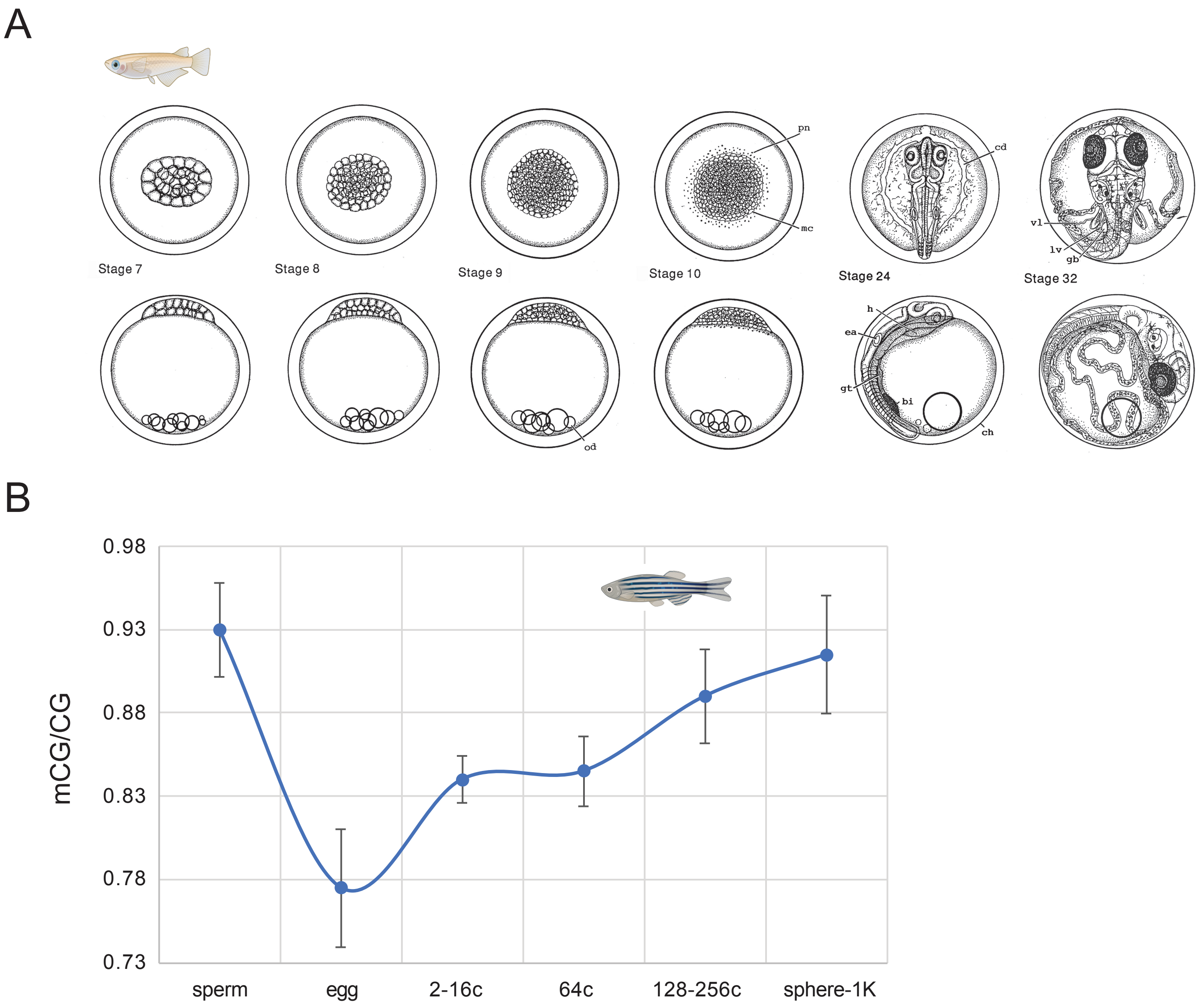


**Supplementary Figure S1. DNA methylome dynamics during early teleost development. (A)** Stages of medaka development used in this study: Stage 7 (32 cells; 3.5hpf), Stage 8 (early morula; 4.5hpf), Stage 9 (late morula; 5.25hpf), Stage 10 (early blastula;6.5hpf), Stage 24 (16 somite; 44hpf), Stage 32 (somite completion; 75hpf). Illustrations from Iwamatsu, 2004 [34]. **(B)** Average DNA methylation levels during zebrafish development. Data correspond to WGBS datasets from [13,14].

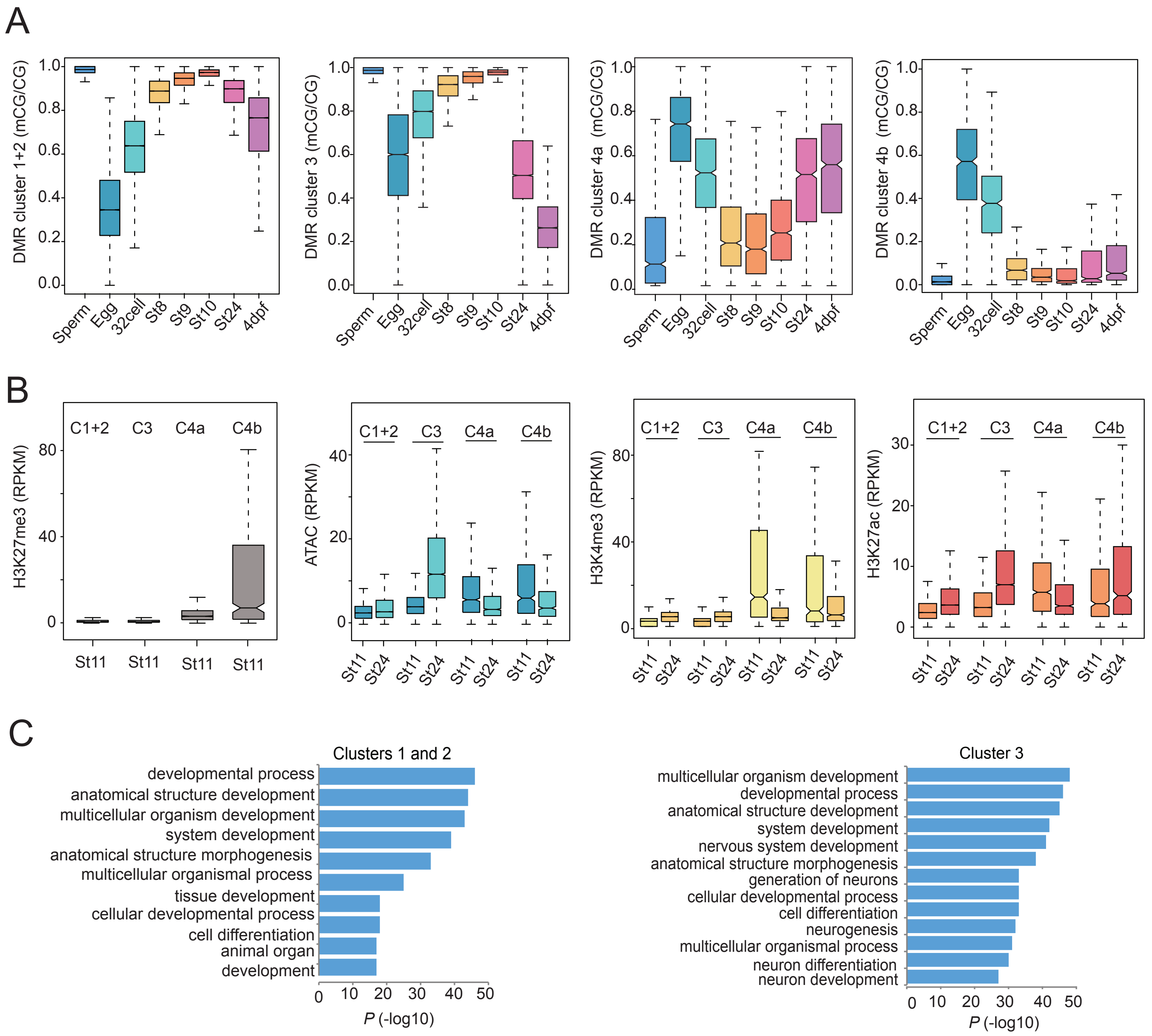


**Supplementary Figure S2. Epigenome remodelling during medaka embryo development. (A)** DNA methylation levels (mCG/CG) during medaka embryo development in differentially methylated regions (DMRs) (cluster 1+2, cluster 3, cluster 4a, cluster4b). The boxplots show the median (centre line) and the first and third quartiles (Q1 and Q3; box limits), and the whiskers extend to the last point within 1.5X the interquartile range below and above Q1 and Q3, respectively. **(B)** ATAC-seq signal and histone modification levels (RPKM) during medaka embryo development (st11, st24) in developmental DMRs (cluster 1+2, cluster 3, cluster 4a, cluster4b). **(C)** Gene Ontology Enrichment analysis on genes which lie withing 5kb of medaka developmental DMRs (cluster 1+2; left, cluster 3; right)

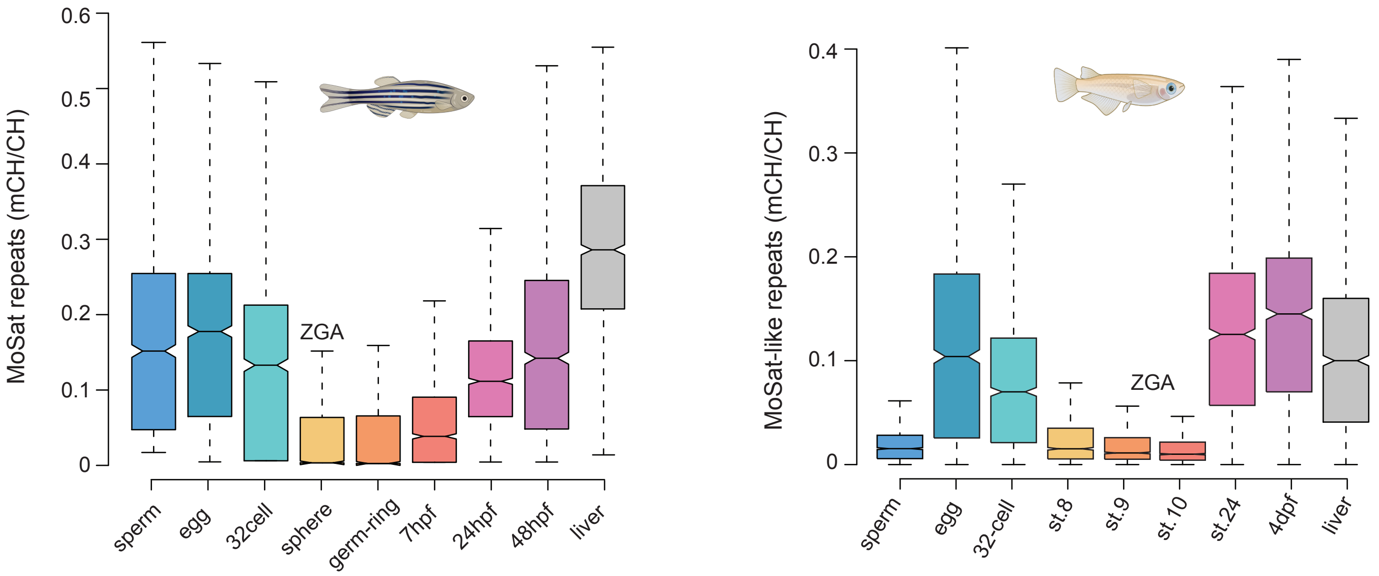


**Supplementary Figure S3. Non-canonical DNA methylation (mCH) reprogramming at mosaic satellite repeats (MoSat) in zebrafish and medaka.** Distribution of CH methylation (mCH/CH) levels at ATGCTA - containing mosaic satellite repeats during zebrafish (left panel) [33] and medaka (right panel) development. The boxplots show the median (centre line) and the first and third quartiles (Q1 and Q3; box limits), and the whiskers extend to the last point within 1.5X the interquartile range below and above Q1 and Q3, respectively.

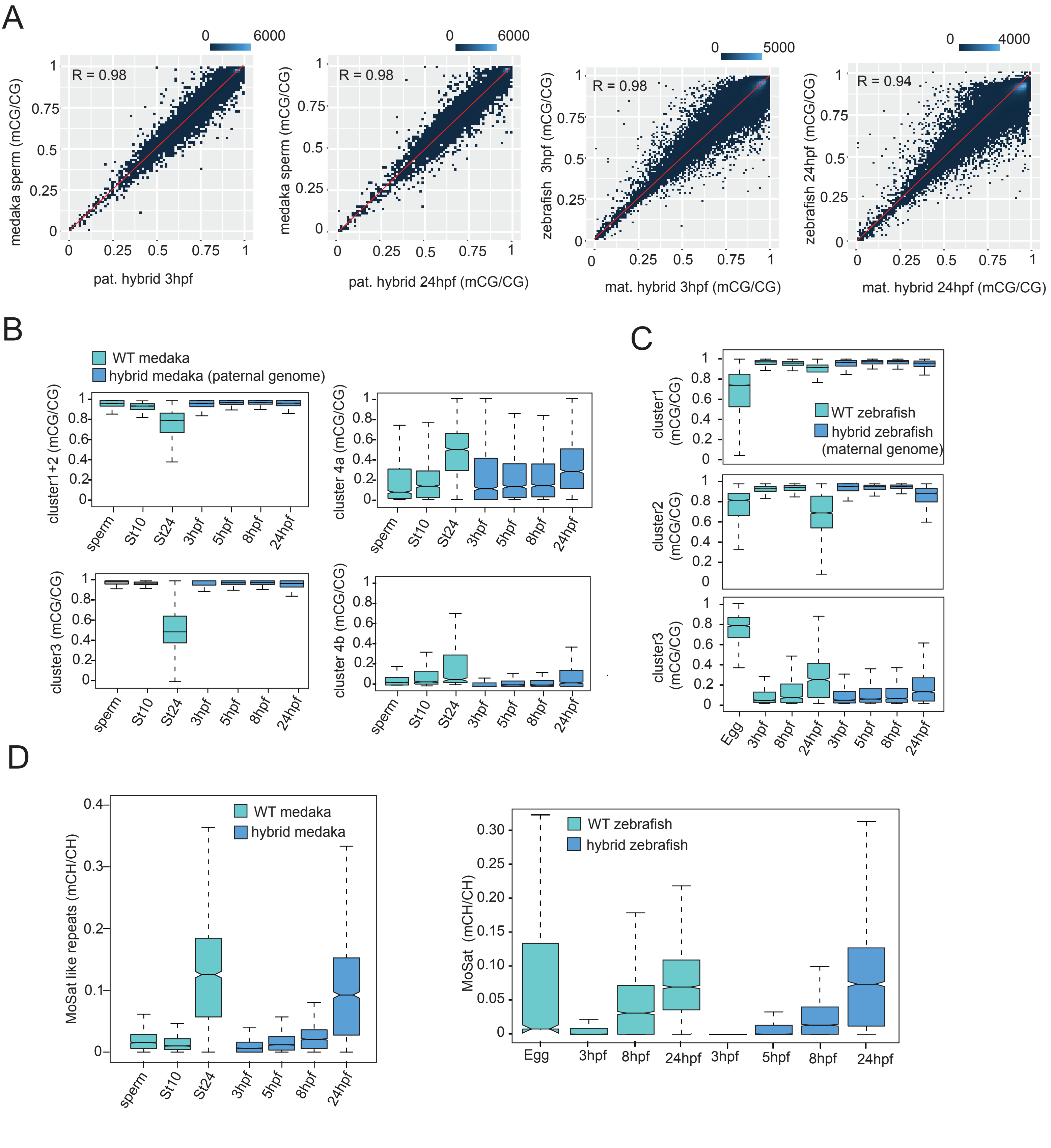


**Supplementary Figure S4. DNA methylation dynamics during medaka-zebrafish hybrid embryo development.** **(A)** Correlation between DNA methylation levels (mCG/CG; 10 kb non-overlapping genomic bins) in medaka sperm vs the paternal medaka methylome in 3hpf and 24hpf medaka-zebrafish hybrid embryos (right); and between 3hpf and 24hpf zebrafish verse the maternal zebrafish methylome in 2hpf and 24hpf medaka-zebrafish hybrid **(B)** DNA methylation levels (mCG/CG) in medaka developmental DMRs in wt medaka and medaka-zebrafish hybrids. The boxplots show the median (centre line) and the first and third quartiles (Q1 and Q3; box limits), and the whiskers extend to the last point within 1.5X the interquartile range below and above Q1 and Q3, respectively. **(C)** DNA methylation levels (mCG/CG) in zebrafish developmental DMRs in wt zebrafish and medaka-zebrafish hybrids. **(D)** CH methylation (mCH/CH) levels at ATGCTA - containing mosaic satellite in the paternal medaka hybrid methylome and wt medaka (left), and the maternal zebrafish hybrid methylome and wt zebrafish (right).
